# Characterization of Endogenous and Epitope-Tagged Ubiquitin Expression and Ubiquitination Patterns at *Drosophila* Neuromuscular Junctions

**DOI:** 10.1101/2025.11.11.687520

**Authors:** Sayantani Kala-Bhattacharjee, Sedonia Davis, Ben Sheline, Craig Schayot, Xiaolin Tian, Chunlai Wu

## Abstract

Ubiquitination is a highly regulated post-translational modification that controls protein stability, activity, localization, and interactions. Disruption of this process contributes to a broad range of human diseases, including cancer, neurodegeneration, and immune dysfunction. The genetic model organism *Drosophila melanogaster* offers powerful tools to study ubiquitination *in vivo*, where versatile systems such as GAL4/UAS, Split-GAL4, GeneSwitch, and MARCM can be combined with imaging and biochemical approaches to achieve precise spatial and temporal analyses. To fully exploit these advantages, experimental designs often require epitope-tagged UAS-Ubiquitin transgenes—both modified and unmodified—that accurately recapitulate the behavior of endogenous Ubiquitin. Here, we report the generation and validation of a panel of epitope-tagged UAS-Ubiquitin transgenes. We first characterize the endogenous ubiquitination pattern at larval neuromuscular junctions (NMJs), then compare that with transgenically expressed Ubiquitin in muscle. Our analysis showed that transgenic His-Myc-, HA-, and His-Biotin-tagged Ubiquitin proteins accurately reproduce endogenous postsynaptic ubiquitination patterns. These validated transgenes provide versatile resources for investigating ubiquitination dynamics across diverse cellular and developmental contexts.

## Introduction

Ubiquitination is a highly conserved post-translational modification that regulates a vast array of cellular processes, including protein turnover, signal transduction, DNA repair, and synaptic plasticity[1–6]. This process is mediated by the covalent attachment of ubiquitin, a small regulatory protein, to substrate proteins in diverse configurations such as mono-ubiquitination or poly-ubiquitin chains of different linkage types. These distinct ubiquitin architectures encode versatile molecular signals that govern substrate stability, localization, and activity. Given its central regulatory role, dysregulation of ubiquitination is implicated in numerous human diseases, including cancer, neurodegeneration, and immune disorders[7,8].

The fruit fly *Drosophila melanogaster* has long served as a powerful genetic system for elucidating fundamental mechanisms of ubiquitination *in vivo*. Its rich genetic toolkit—featuring the GAL4/UAS[9], Split-GAL4[10], GeneSwitch[11], and MARCM[12] systems—enables precise spatial and temporal control of gene expression. When combined with imaging, electrophysiological, and biochemical analyses, these tools allow detailed dissection of ubiquitin-dependent pathways within specific cell types and developmental stages.

Despite these advantages, there remains a lack of versatile resources for directly manipulating ubiquitin itself in *Drosophila*. Existing reagents largely focus on components of the ubiquitination machinery, such as E2 conjugating enzymes or E3 ligases, while tools that permit controlled expression and tracking of ubiquitin remain limited. In particular, UAS-driven, epitope-tagged ubiquitin transgenes that faithfully mimic endogenous ubiquitin behavior are essential for visualizing and biochemically characterizing ubiquitination events in specific tissues.

In this study, we describe the generation and validation of a series of epitope-tagged UAS-Ubiquitin transgenes designed for tissue-specific expression in *Drosophila*. Comparative analyses of these constructs at the neuromuscular junction (NMJ) reveal that His-Myc–tagged, HA-tagged, and His-Biotin-tagged ubiquitin uniquely recapitulate the postsynaptic ubiquitination patterns observed with endogenous ubiquitin. These validated reagents provide a robust platform for investigating ubiquitination dynamics with high spatial and temporal precision, greatly expanding the experimental capacity for studying protein regulation and signaling in neuronal and non-neuronal contexts alike.

## Results

### Characterization of endogenous ubiquitination patterns at *Drosophila* larval NMJs and ventral nerve cord

After evaluating a panel of anti-ubiquitin antibodies, we identified the mouse monoclonal anti–mono- and poly-ubiquitinylated conjugates antibody (FK2) as the only reagent that yielded reliable immunostaining results. In wild-type larval body wall muscles, FK2-positive signals were detected throughout the muscle but were particularly enriched at the postsynaptic density (PSD) of the neuromuscular junctions (NMJs). Co-staining with the presynaptic vesicle marker DvGlut (Drosophila Vesicular Glutamate Transporter) revealed that ubiquitinylated conjugates encircle the presynaptic terminals, forming an embedded ring-like structure (Fig. 1A). A similar pattern was observed when FK2 staining was combined with HRP, a presynaptic membrane marker (Fig. 1B).

**Figure 1.**
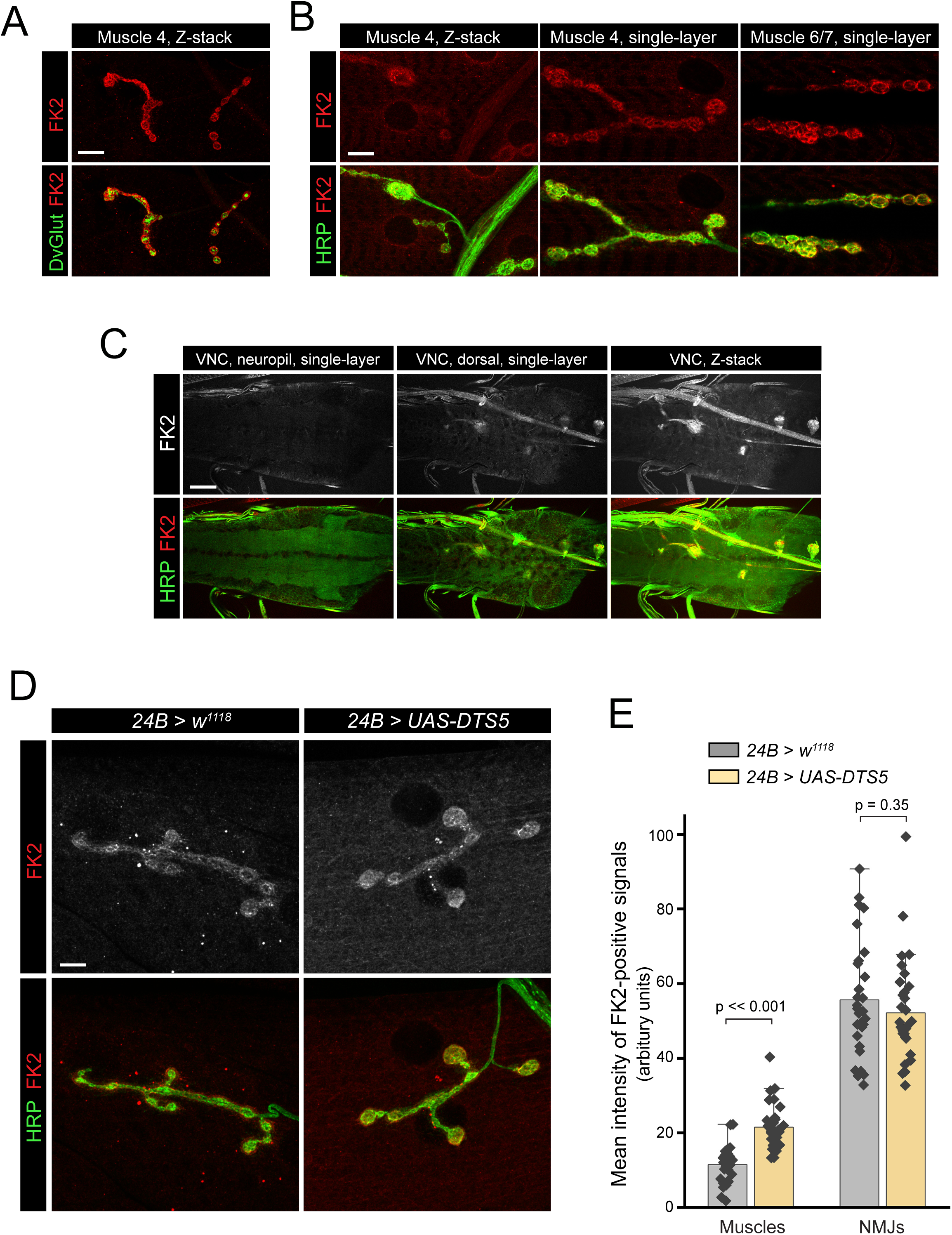
Expression pattern of endogenous ubiquitination at larval NMJs and ventral nerve cord (VNC). (**A-C**) Representative confocal images showing ubiquitination patterns at larval neuromuscular junctions (NMJs; A-B) and the ventral nerve cord (VNC; C), immunostained with anti-FK2 (ubiquitin conjugates) and anti-HRP (neuronal membrane marker). Scale bars, for A: 12 μm; for B: 8 μm; for C: 40 μm. (**D**) Representative confocal images of muscle 4 NMJs from control larvae and larvae expressing UAS-DTS5 under the 24B-Gal4 driver, stained with FK2 and HRP. Scale bar: 8 μm. (**E**) Quantification of mean FK2-positive signal intensity at larval muscles and NMJs comparing control and *24B>UAS-DTS5* larvae, illustrating the effect of muscle-specific DTS5 overexpression on ubiquitination levels.

The FK2-labeled signals appeared either diffuse or punctate within both muscle fibers and NMJ regions, indicating heterogeneity in the distribution of ubiquitinylated conjugates. These observations suggest that ubiquitinylated conjugates are predominantly localized at the postsynaptic sites of NMJs. In contrast, in the larval ventral nerve cord, FK2 staining intensity was relatively weak in both the synapse-rich neuropil and the surrounding cell body layer (Fig. 1C).

To assess the sensitivity of FK2 staining to proteasomal inhibition, we expressed UAS-DTS5, which encodes a dominant-negative mutant of the proteasomal β6 catalytic subunit[13], in muscle using the 24B-Gal4 driver. Comparison of FK2 immunoreactivity between control and DTS5-expressing larvae revealed a significant accumulation of ubiquitinylated conjugates in the muscle cytoplasm, whereas signal intensity at the NMJs remained unchanged (Fig. 1D-E). These findings indicate that ubiquitinylated conjugates at NMJs are likely enriched in mono-ubiquitinated species that are not subject to proteasomal degradation. Moreover, these results validate that the FK2 antibody specifically recognizes ubiquitinylated proteins in *Drosophila* tissues.

### Generation of wild type and mutated Ubiquitin transgenes under UAS-promoter

Ubiquitin can be covalently attached to one or multiple lysine residues of a target protein, resulting in mono- or polyubiquitination[14]. The fate of a ubiquitinated protein depends on the type and topology of the ubiquitin modification[15]. Ubiquitin itself contains eight amino groups (M1, K6, K11, K27, K29, K33, K48, and K63) capable of forming a wide array of homotypic or heterotypic (branched) polyubiquitin chains, thereby providing remarkable regulatory diversity. Among these, K48-linked polyubiquitin chains are best known for targeting substrates for degradation via the 26S proteasome. In contrast, monoubiquitination often alters protein–protein interactions, subcellular localization, or folding stability rather than promoting degradation [16–18].

To facilitate *in vivo* analysis of ubiquitination mechanisms, we generated two ubiquitin mutants—Ub[K48R] and Ub[K0]—in addition to wild-type ubiquitin. The Ub[K48R] mutant specifically blocks chain elongation at lysine 48, while Ub[K0], a lysine-less variant with all seven lysines replaced by arginines, can be conjugated to substrates but cannot serve as a polymerization site, thereby preventing polyubiquitin chain formation. Wild-type and mutant ubiquitins were subcloned into pUAST-His-Myc, pUAST-HA, pUAST-His-Biotin, and pUAST-NTAP vectors to generate a panel of epitope-tagged UAS-Ubiquitin transgenes (Table 1).

**Table 1.**
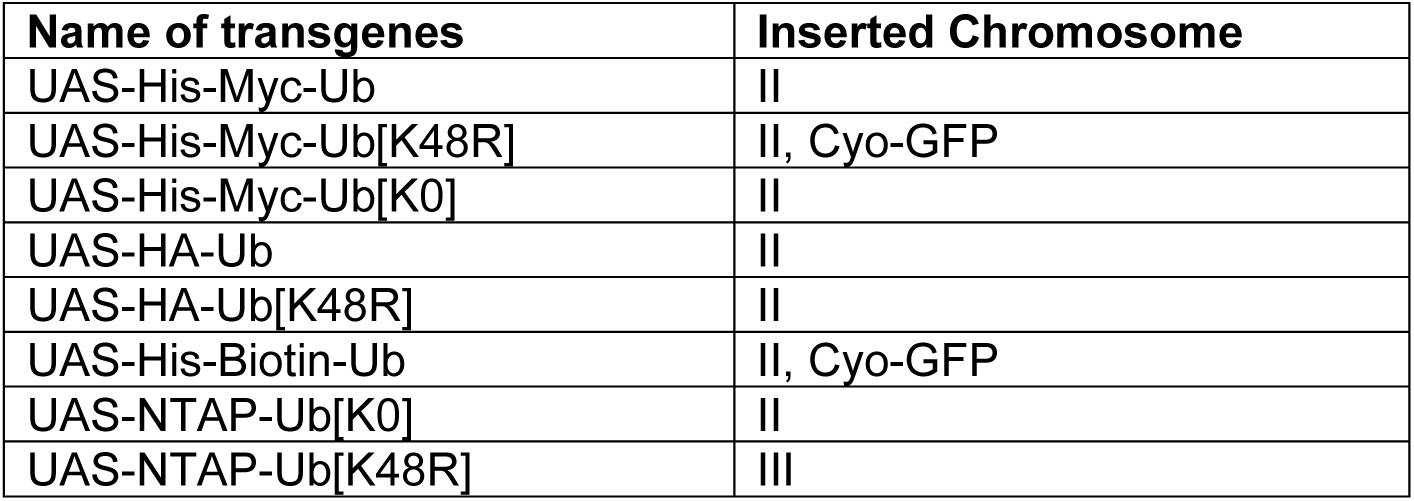
UAS-epitope-tagged transgenes generated.

### Validation of UAS-Ubiquitin transgenes in immunocytochemistry

#### UAS-His-Myc-Ubiquitin

When His-Myc–tagged wild-type ubiquitin (HM-Ub), Ub[K48R], or Ub[K0] were expressed in larval muscles, all three proteins showed strong enrichment at the postsynaptic density (PSD), closely resembling the endogenous ubiquitination pattern at NMJs (Fig. 2A). Co-immunostaining with the FK2 antibody revealed that PSD-enriched HM-Ub, HM-Ub[K48R], and HM-Ub[K0] were all recognized by FK2, indicating that each tagged ubiquitin species participates in forming ubiquitinylated conjugates at the PSD.

**Figure 2.**
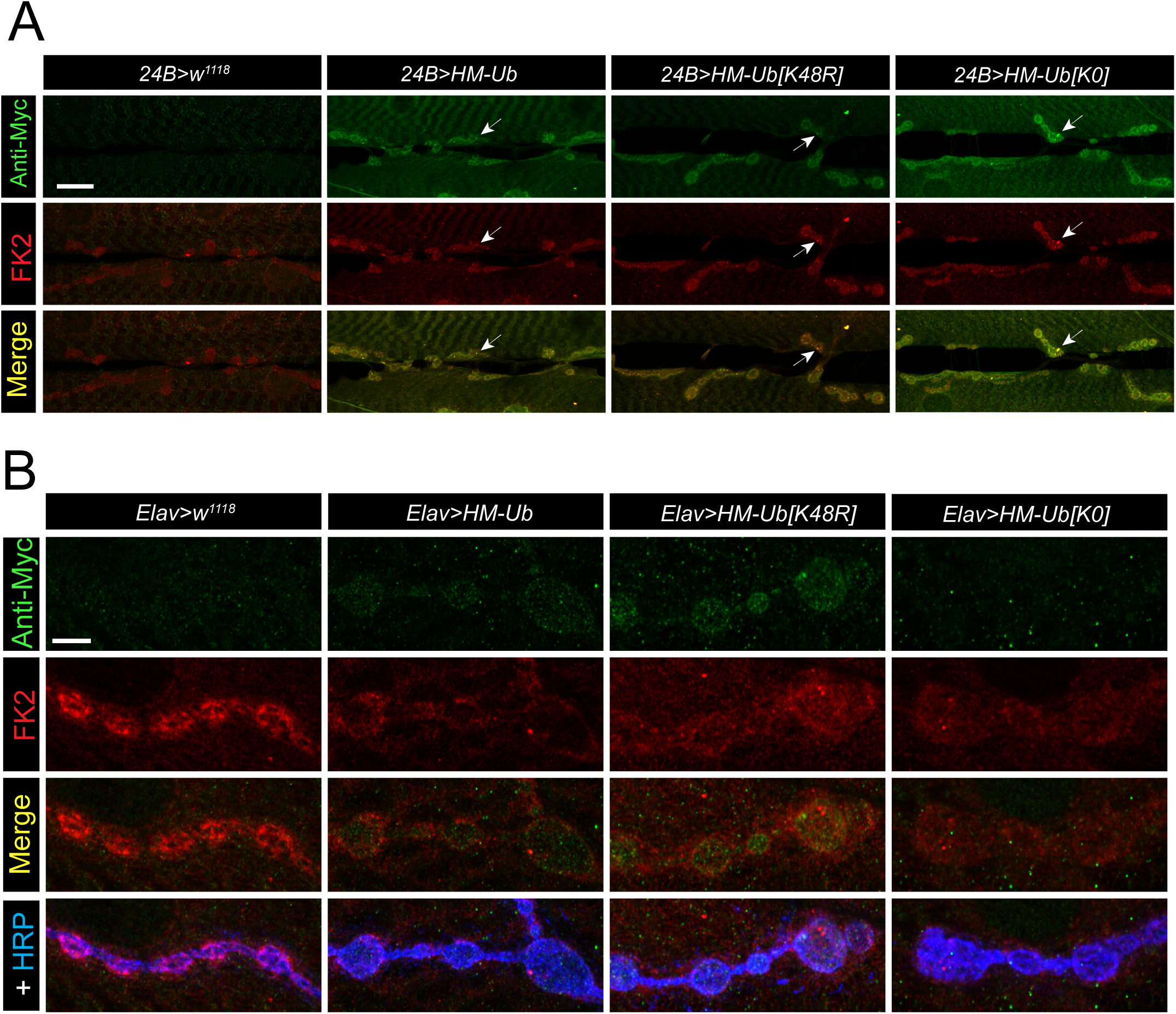
Expression pattern of muscle and neuronal expressed HM-Ub, HM-Ub [K48R], and HM-Ub [K0] transgenes. (**A**) Representative single-layer confocal images show ubiquitination patterns at larval muscle 6/7 from control larvae (*24B>w^1118^*) and from larvae expressing *UAS-HM-Ub*, *UAS-HM-Ub[K48R]*, and *UAS-HM-Ub[K0]* in muscles under the control of 24B-Gal4. Samples were immunostained with anti-Myc and anti-FK2 antibodies. Arrows indicate puncta positive for both Myc and FK2 signals. Scale bar: 20 μm. (**B**) Representative single-layer confocal images show ubiquitination patterns at the tips of larval muscle 4 from control larvae (*Elav>w^1118^*) and from larvae expressing *UAS-HM-Ub*, *UAS-HM-Ub[K48R]*, and *UAS-HM-Ub[K0]* in neurons under the control of Elav-Gal4. Samples were immunostained with anti-Myc, anti-FK2, and anti-HRP antibodies. Note that HM-Ub[K0] lacks detectable presynaptic staining. No overlap was observed between Myc-positive puncta and FK2 signals in HM-Ub expressing NMJs, whereas mild colocalization was detected in NMJs expressing HM-Ub[K48R]. Scale bar: 5 μm.

In contrast, when the same transgenes were expressed in neurons, only wild-type HM-Ub and HM-Ub[K48R]—but not HM-Ub[K0]—were detected as fine puncta at NMJs. Single-layer confocal scans of Myc- and FK2-positive signals revealed no overlap between HM-Ub and FK2, and only minimal colocalization for HM-Ub[K48R] (Fig. 2B). These observations suggest that most presynaptically localized HM-Ub molecules exist in an unconjugated form and that the FK2 antibody selectively recognizes mono- and poly-ubiquitinated conjugates rather than free ubiquitin.

In the ventral nerve cord (VNC), HM-Ub[K48R] exhibited the highest expression level, followed by HM-Ub and HM-Ub[K0] (Fig. S1). Double labeling with FK2 showed that neither the cell bodies nor the neuropil regions expressing HM-tagged Ubs were FK2 positive, further supporting that these Myc-labeled signals represent unconjugated ubiquitin species. Notably, HM-Ub[K0] displayed the lowest expression level in the VNC but the highest relative enrichment at the NMJ PSD, suggesting that lysine-less ubiquitin preferentially participates in mono-ubiquitination at the postsynaptic site.

#### UAS-HA-Ubiquitin

The HA-tagged Ub and Ub[K48R] variants exhibited expression patterns similar to their His-Myc–tagged counterparts. When expressed in muscles, both HA-Ub and HA-Ub[K48R] were highly enriched at the postsynaptic density (PSD) of the NMJs and were recognized by the FK2 antibody (Fig. 3A), indicating that both form ubiquitinated conjugates. In contrast, when expressed neuronally, HA-Ub was primarily localized to neuronal cell bodies within the ventral nerve cord (VNC) (Fig. S2). At the NMJs, only weak punctate HA-Ub and HA-Ub[K48R] signals were detected at the terminals of muscle 6/7, with little to no staining observed at other NMJs (Fig. 3B). Double immunostaining with anti-HA and anti-FK2 antibodies revealed that FK2 did not recognize the HA-positive staining at neuronal cell bodies at VNC, nor the fine HA-positive puncta at the tip of muscle 6/7, suggesting that they predominantly represent soluble HA-Ub or HA-Ub[K48R] monomers rather than ubiquitinated conjugates.

**Figure 3.**
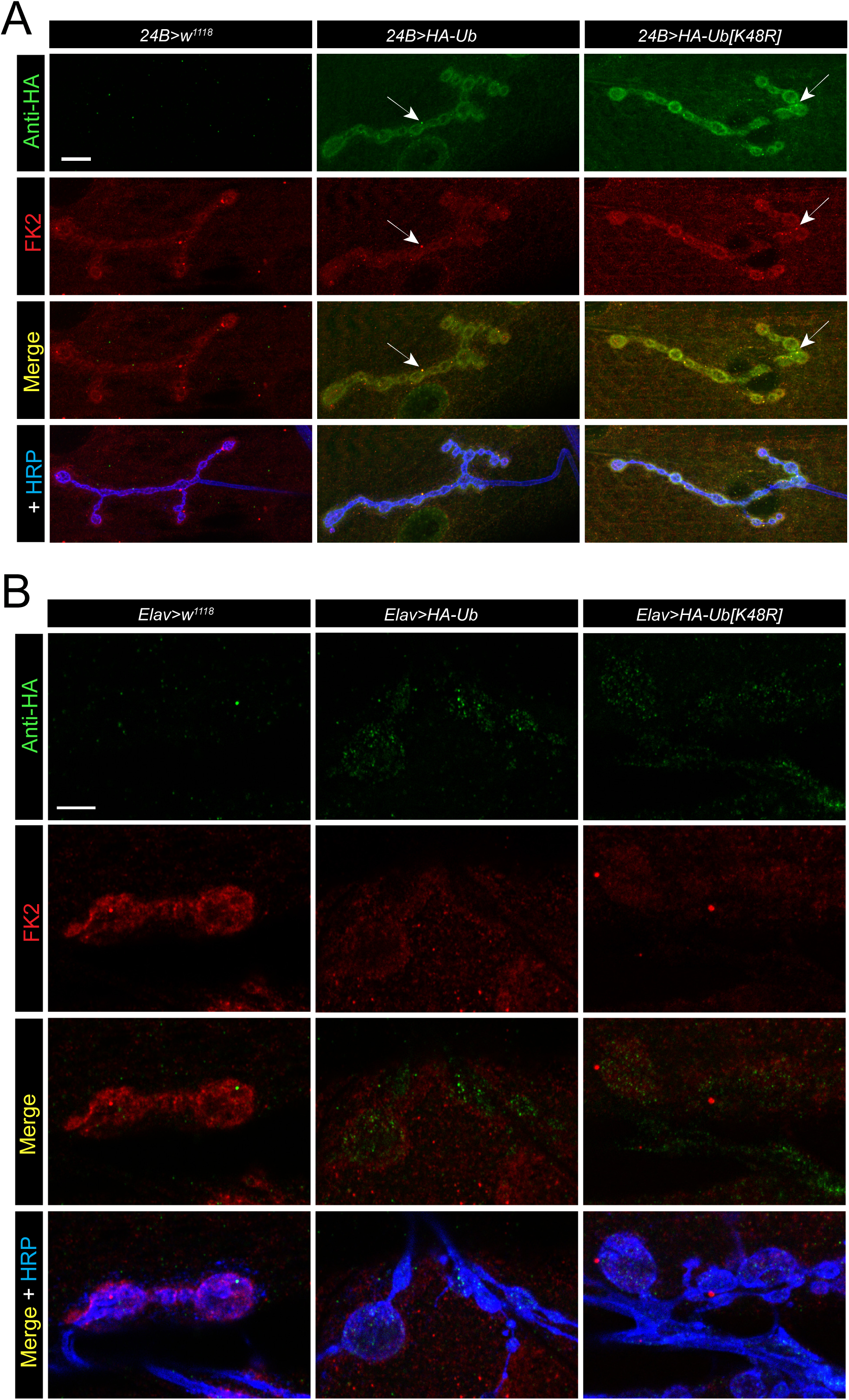
Expression pattern of muscle- and neuron-expressed HA-Ub and HA-Ub [K48R] transgenes at NMJs. (**A**) Representative single-layer confocal images show ubiquitination patterns at larval muscle 4 from control larvae (*24B>w^1118^*) and from larvae expressing HA-Ub and HA-Ub[K48R] in muscles under the control of 24B-Gal4. Samples were immunostained with anti-HA, anti-FK2, and anti-HRP antibodies. Arrows indicate puncta positive for both HA and FK2 signals. Scale bar: 10 μm. (**B**) Representative single-layer confocal images show ubiquitination patterns at the tips of larval muscle 6/7 from control larvae (*Elav>w^1118^*) and from larvae expressing HA-Ub and HA-Ub[K48R] in neurons under the control of Elav-Gal4. Samples were immunostained with anti-HA, anti-FK2, and anti-HRP antibodies. Note that there is almost no overlap of the HA-positive puncta with the FK2 signals. Scale bar: 4 μm.

#### UAS-His-Biotin-Ubiquitin

Muscle-expressed His-Biotin–tagged Ubiquitin is highly enriched at the postsynaptic density of neuromuscular junctions (NMJs), closely resembling the endogenous ubiquitination pattern (Fig. 4). The HB-Ubiquitin–positive signal is also recognized by the FK2 antibody, indicating that HB-Ubiquitin forms mono- or polyubiquityl conjugates at the postsynaptic sites (Fig. 4). In contrast, neuronally expressed HB-Ubiquitin is predominantly localized in cell bodies, the synapse-rich neuropil of the ventral nerve cord, and motor neuron axons (Fig. S3). Although mild staining is observed at presynaptic terminals, its distribution—characterized by a gradient extending from distal axons—suggests that these signals likely represent passive diffusion from axons rather than active synaptic localization. Notably, none of the neuronal HB-Ubiquitin signals are detected by the FK2 antibody, indicating that the Biotin-positive signals in the nervous system predominantly correspond to soluble HB-Ubiquitin monomers rather than ubiquitylated conjugates.

**Figure 4.**
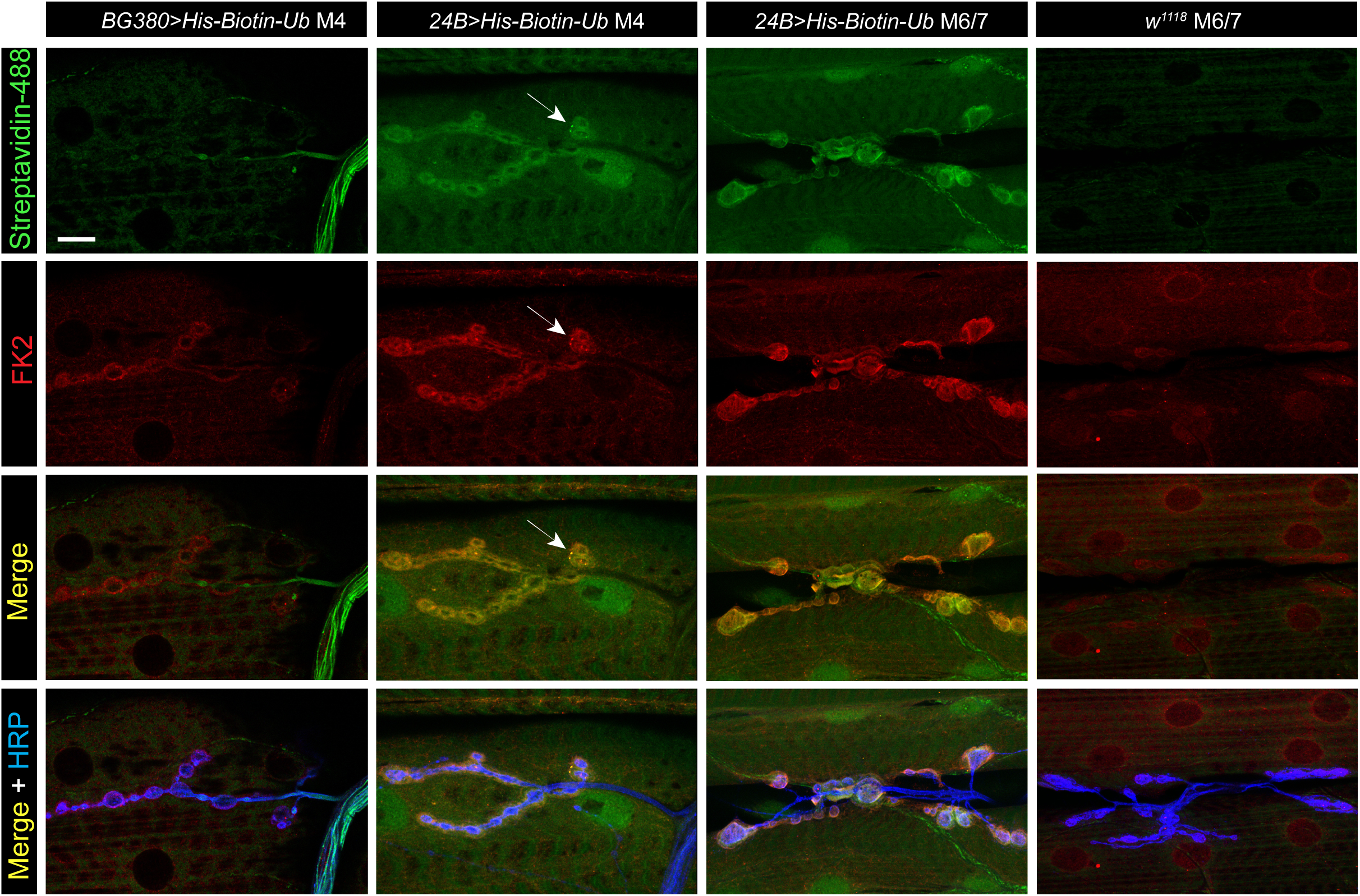
Expression pattern of muscle- and neuron-expressed His-Biotin-Ubiquitin transgene at NMJs *UAS-His-Biotin-Ubiquitin* was expressed in neurons and muscles using BG380-Gal4 and 24B-Gal4 drivers, respectively. Representative single-layer confocal images show ubiquitination patterns at larval muscle 4 (M4) and muscle 6/7 (M6/7) NMJs from wild-type larvae and from larvae expressing HB-Ub in neurons or muscles. Note the increased FK2 signal intensity at NMJs in 24B-driven HB-Ub larvae compared to wild-type and BG380-driven HB-Ub larvae. Arrows indicate puncta that are positive for both streptavidin and FK2 staining. Scale bar: 10 μm.

#### UAS-NTAP-Ubiquitin

Building on our previous success using the tandem affinity purification (TAP) system to identify protein–protein interactions[19], we next tested whether a TAP tag could be fused to ubiquitin to enable two-step purification of ubiquitinated proteins in a tissue-specific manner. Neuronal expression of NTAP-Ub[K48R] and NTAP-Ub[K0] in the larval brain resulted in clear labeling within the VNC (Fig. S4 A). In contrast, when NTAP-Ub[K48R] and NTAP-Ub[K0] were individually expressed in larval muscles, TAP-immunoreactivity was absent from the postsynaptic density, differing from the distribution pattern observed for endogenous ubiquitin (Fig. S4 B). This suggests that NTAP-Ub was unable to conjugate efficiently to postsynaptic substrates. A plausible explanation is that the relatively large (∼26 kDa) TAP epitope may disrupt proper folding of the ubiquitin molecule or hinder its recognition by E2/E3 enzymes, thereby impairing conjugation. In contrast, smaller tags such as His-Myc, HA, and His-Biotin do not appear to interfere with ubiquitin’s normal physiological functions.

## Discussion

Post-synaptic proteins are highly ubiquitinated[5]. Previous studies have shown that synaptic activity enhances ubiquitin conjugation of PSD proteins in cultured primary neurons, and that proteins within the postsynaptic apparatus undergo rapid turnover mediated by the ubiquitin–proteasome system (UPS)[20]. Such a dynamic protein turnover at the postsynaptic density is crucial for normal postsynaptic function and plasticity[5]. Consistent with this, we found that endogenous ubiquitination is primarily localized to postsynaptic sites. However, our DTS5 experiments indicate that the prominent ubiquitination events at larval NMJs are likely mono-ubiquitination, which may play an important role in postsynaptic receptor regulation. Recent proteomic analyses have identified some of the ubiquitinated proteins in fly neuronal tissue, revealing 48 newly ubiquitinated proteins, many of which are critical for synaptogenesis[21]. Collectively, these findings suggest that *Drosophila* NMJs provide a powerful model for studying ubiquitin-dependent pathways that regulate synaptic structure and function.

Under physiological conditions, ubiquitin is not synthesized as a free monomer but rather as a linear fusion precursor. These precursors are produced either as polyubiquitin chains encoded by stress-inducible genes containing tandem repeats of the ubiquitin coding sequence, or as ubiquitin–ribosomal subunit fusion proteins. In both cases, the precursor proteins are post-translationally processed by deubiquitinating enzymes to release functional monomeric ubiquitin. Based on this understanding, transgenic strategies expressing ubiquitin precursors have been developed in both *Drosophila* and mouse to study *in vivo* ubiquitination. For example, the *Drosophila* UAS-(bioUb)₆-BirA transgene encodes six tandem ubiquitin units carrying a BirA recognition motif followed by BirA[21], whereas the murine 6xHis-Ub-GFP transgene fuses His-tagged ubiquitin with GFP[22]. In both systems, the tagged ubiquitin monomers and their fusion partners are presumably released through the action of endogenous deubiquitinating enzymes, and both biotinylated and His-tagged ubiquitin were shown to conjugate efficiently to cellular substrates. In contrast, our study shows that precursor processing is not essential for functional ubiquitin conjugation in *Drosophila*. The epitope-tagged ubiquitins we generated are expressed as monomers and form conjugates that closely match the patterns of endogenous ubiquitination at larval muscles. This suggests that monomeric, properly folded ubiquitin— even when epitope-tagged—can fully engage in the normal ubiquitination machinery. Importantly, this approach provides a simpler and more biologically faithful system for studying ubiquitination *in vivo*. By expressing ubiquitin directly as a monomer, our transgenes avoid potential artifacts from polyubiquitin fusion constructs and offer a flexible, reliable toolkit for analyzing ubiquitin dynamics, substrate specificity, and proteostasis across different tissues.

A ubiquitin variant carrying a noncleavable C-terminal substitution (UbG76V) has been fused to GFP to monitor proteasomal activity. The UbG76V-GFP fusion protein is short-lived and rapidly degraded by the proteasome under normal conditions, whereas impairment of the ubiquitin–proteasome system (UPS) leads to accumulation of GFP fluorescence[23]. Thus, UbG76V-GFP transgenes targeted to specific subcellular compartments can serve as reporters for tissue-specific UPS function. However, expression of UbG76V-GFP fused to synaptobrevin-2 (Syb-2) in mouse neurons caused adult-onset degeneration of motor neuron terminals, suggesting that the localization signal critically determines the toxicity of the fusion protein[24]. In contrast, muscle-or neuron-specific expression of our epitope-tagged ubiquitin constructs produced no detectable morphological abnormalities at the NMJ. Moreover, UPS impairment increased HM-Ub accumulation in a cell-type–specific manner. Together, these findings indicate that our tagged ubiquitin reporters provide a reliable and non-toxic tool for monitoring UPS activity *in vivo*.

The three epitope tags used to generate the Ubiquitin transgenes—His-Myc (HM), HA, and His-Biotin (HB)—are small and minimally invasive. Our data indicate that all three tagged forms effectively mimic the behavior of endogenous ubiquitin, including conjugation to target proteins. These epitope-tagged ubiquitins offer high flexibility, sensitivity, and reliability for visualizing tissue-specific ubiquitination through immunostaining and imaging. In addition to their imaging utility, the His tag present in the HM- and HB-Ub constructs enables metal-chelate affinity purification, while the dual His-Biotin tag allows tandem affinity purification using sequential Ni²⁺-NTA and streptavidin resins. This two-step approach enhances purification specificity and reduces background contamination. Collectively, these transgenes provide a versatile toolkit for investigating ubiquitination at cellular, molecular, and biochemical levels.

Our data suggest that in the *Drosophila* nervous system, particularly within the central nervous system (CNS), overall ubiquitination levels are extremely low and often near the limit of detection. This likely reflects both a relatively low basal rate of ubiquitination and the rapid turnover of ubiquitinated proteins by an active ubiquitin–proteasome system (UPS). To investigate how neuronal ubiquitination dynamics change in response to specific genetic or physiological stimuli, it would be advantageous to co-express temperature-sensitive proteasome mutants such as UAS-DTS5 or UAS-DTS7 alongside UAS-tagged-Ubiquitin transgenes[13,25]. By rearing flies at different temperatures, the degree of UPS inhibition can be precisely modulated, allowing controlled accumulation of ubiquitinated proteins. Such a strategy provides an effective means to capture and analyze ubiquitination events that would otherwise be too transient to detect under normal physiological conditions.

## Materials and Methods

### Drosophila strains

The following strains were used in this study: C155-Gal4 (Elav-Gal4, Bloomington *Drosophila* Stock Center [BDSC] #458), 24B-Gal4 (muscle-specific), *w^1118^* (BDSC #6326), BG380 (neuron specific)[26]. The UAS-DTS5 transgene is a gift from Dr. John Belote. Flies were reared on standard cornmeal-yeast-agar medium at 25°C under a 12 h light/dark cycle.

### Generation of Epitope-tagged Ubiquitin transgenes

Wild type, K48R, K0 ubiquitin cDNA constructs were gifts from Dr. Richard Baer[27]. Wild type and mutated Ub were subcloned into pUAST-His-Myc, pUAST-HA, pUAST-His-Biotin, and pUAST-NTAP vectors to generate the corresponding transformation DNA constructs. These constructs were then injected into fly embryos using standard procedure[28] to generate a panel of epitope-tagged UAS-Ubiquitin transgenes (Table 1). These transgenes will be available via the Bloomington Stock Center for distribution.

### Immunocytochemistry

Larval muscles were dissected in ice-cold phosphate-buffered saline (PBS) and fixed in 4% paraformaldehyde for 20 minutes, except for samples processed for anti-DvGlut staining, which were fixed in cold methanol for 5 minutes. The fixed tissues were immuno-stained following standard procedures. The primary antibodies used were: mouse anti-ubiquitin at 1:1000 (Millipore, Clone FK2, 04–263), rabbit anti-DvGlut at 1:5000[29], rabbit anti-Myc at 1:2000 (Bethyl, A190-105A), rabbit anti-HA at 1:1000 (Cell Signaling Technology, 3724S), goat anti-HRP conjugated with Alexa Fluor 488 or Alexa Fluor 647 (Jackson ImmunoResearch). The Biotin tag was detected using Streptavidin conjugated with Alexa Fluor 488 (Jackson ImmunoResearch). The following Secondary antibodies (Jackson ImmunoResearch) included DyLight 549–conjugated goat anti-mouse IgG (1:1000) and DyLight 488–conjugated goat anti-rabbit IgG (1:1000).

### Imaging and quantification at third instar larvae

Single-layer or Z-stack confocal images were acquired using a Nikon C1 confocal microscope (Melville, NY). Unless otherwise noted, images displayed within the same picture panel were captured under identical gain settings from samples that were fixed and stained in the same dissection dish. Quantification of FK2 mean signal intensity at the muscle regions and NMJs was performed using ImageJ (FIJI) in a blinded manner.

### Statistical analysis

Statistical analyses graph generation were conducted using OriginPro (OriginLab, Northampton, MA, USA). Comparisons between two groups were evaluated using Student’s *t*-test. Bar-overlap plots represent mean ± s.e.m., with all individual data points displayed.

---

**Supplemental Figure 1.**
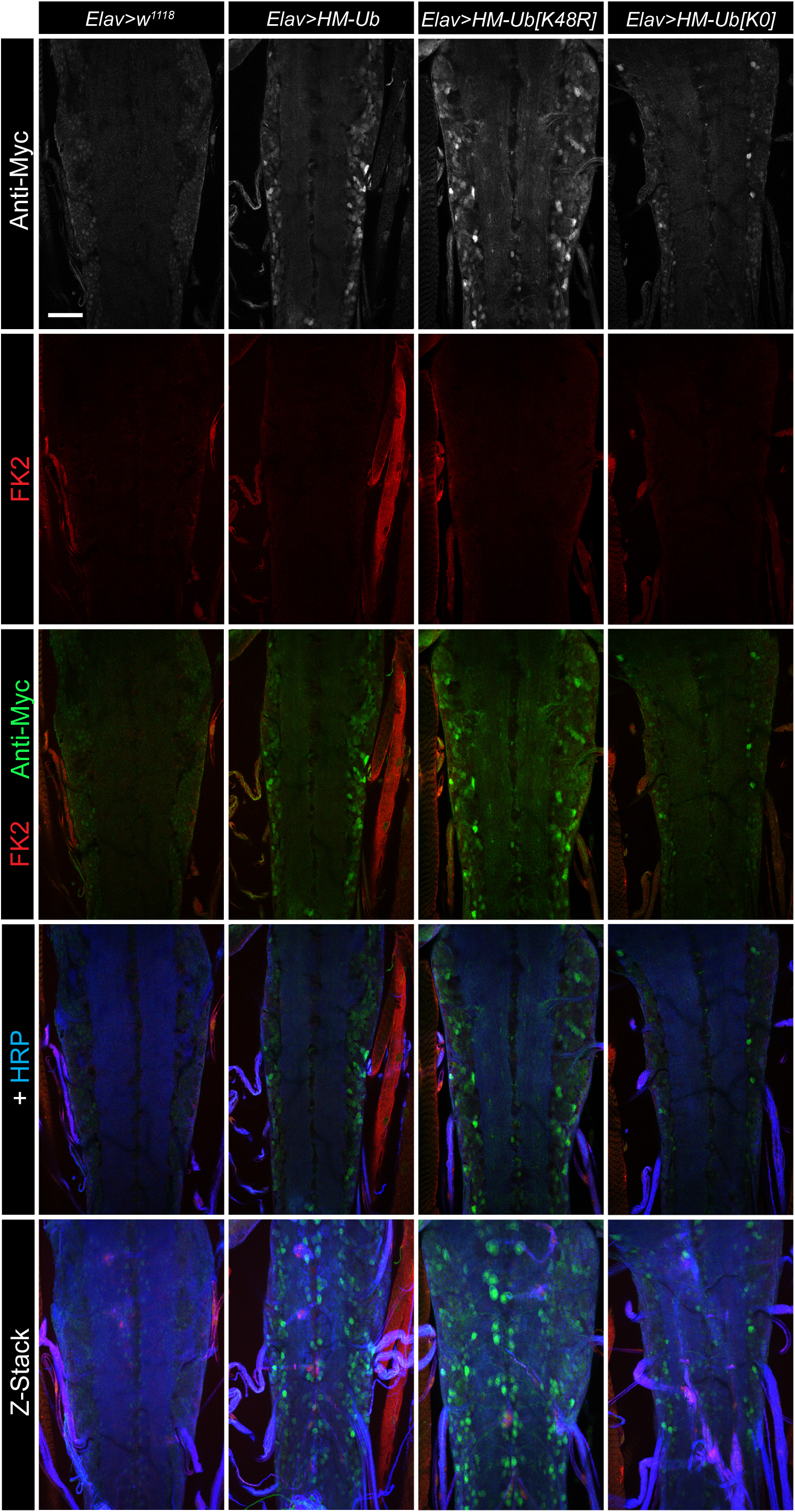
Expression of HM-Ub, HM-Ub [K48R], and HM-Ub [K0] in the larval VNC Representative confocal images of larval ventral nerve cords (VNCs) from control (*Elav>w^1118^*) and flies expressing Elav-driven UAS-HM-Ub, UAS-HM-Ub(K48R), or UAS-HM-Ub(K0) transgenes. Samples were immunostained with anti-Myc, anti-FK2, and anti-HRP antibodies. All images represent single confocal sections focused on the neuropil region of the VNC, except for the bottom row, which shows Z-stack projections of the entire VNC. Scale bar: 35 μm

**Supplemental Figure 2.**
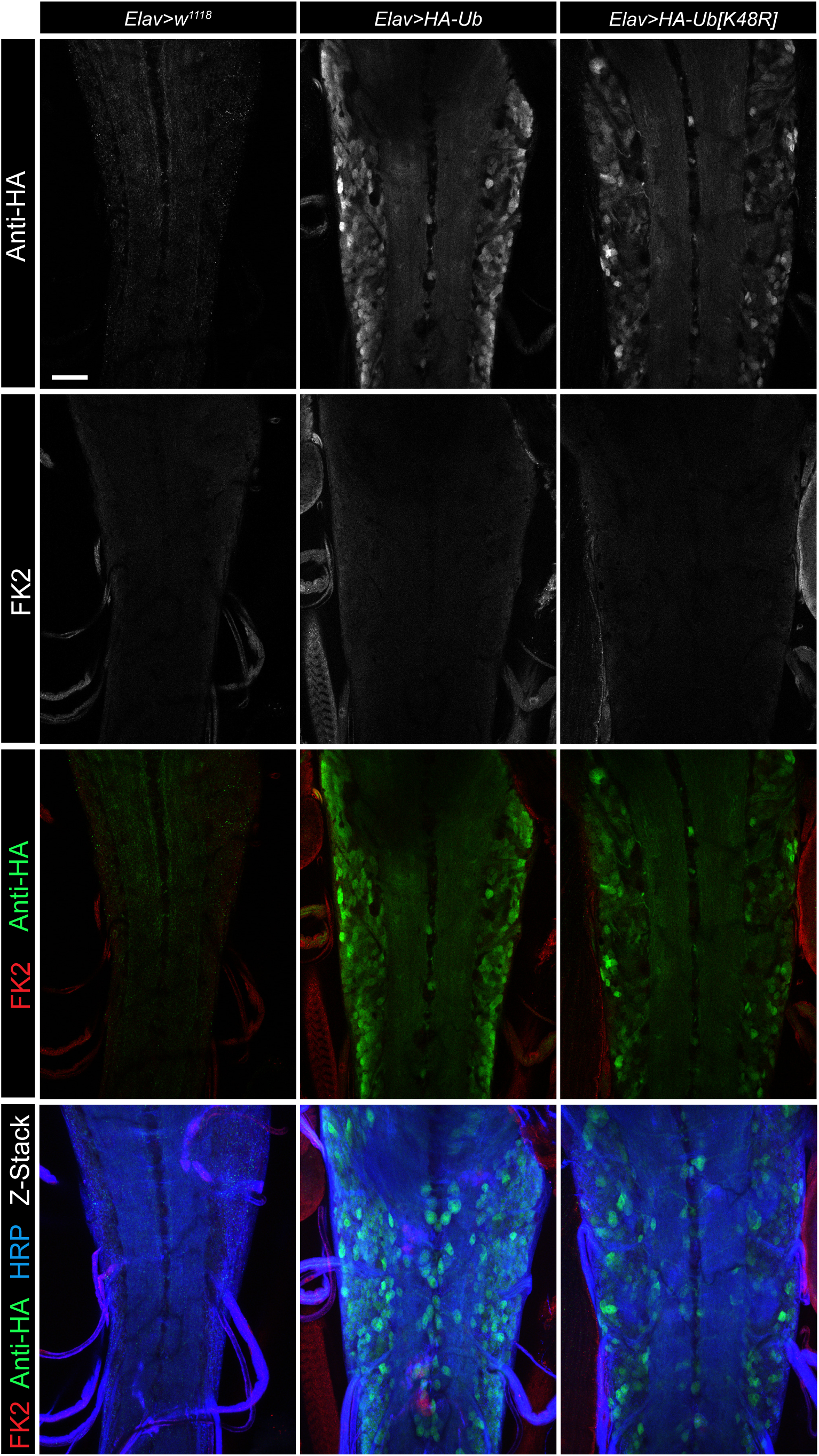
Expression of HA-Ub, and HA-Ub [K48R] in the larval VNC Representative confocal images of larval ventral nerve cords (VNCs) from control (*Elav>w^1118^*) and flies expressing Elav-driven UAS-HA-Ub and UAS-HA-Ub(K48R). Samples were immunostained with anti-HA, anti-FK2, and anti-HRP antibodies. All images represent single confocal sections focused on the neuropil region of the VNC, except for the bottom row, which shows Z-stack projections of the entire VNC. Scale bar: 35 μm.

**Supplemental Figure 3.**
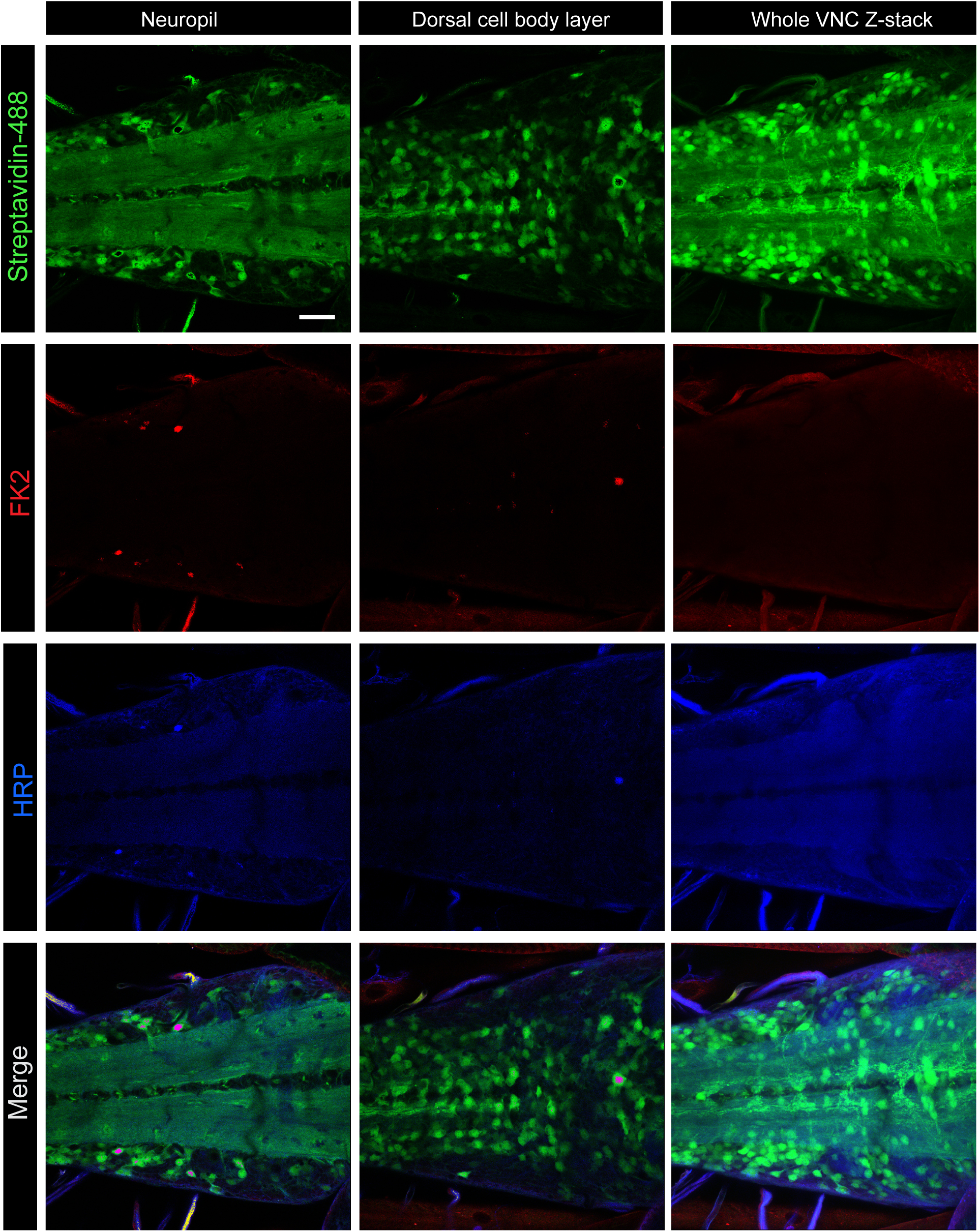
Expression of His-Biotin-Ub in the larval VNC Representative confocal images of larval VNCs from flies expressing *UAS-His-Biotin-Ubiquitin* under the control of the BG380-Gal4 driver. Shown are single confocal sections focused on the neuropil region and the dorsal cell body layer, as well as Z-stack projections encompassing the entire VNC. Scale bar: 20 μm.

**Supplemental Figure 4.**
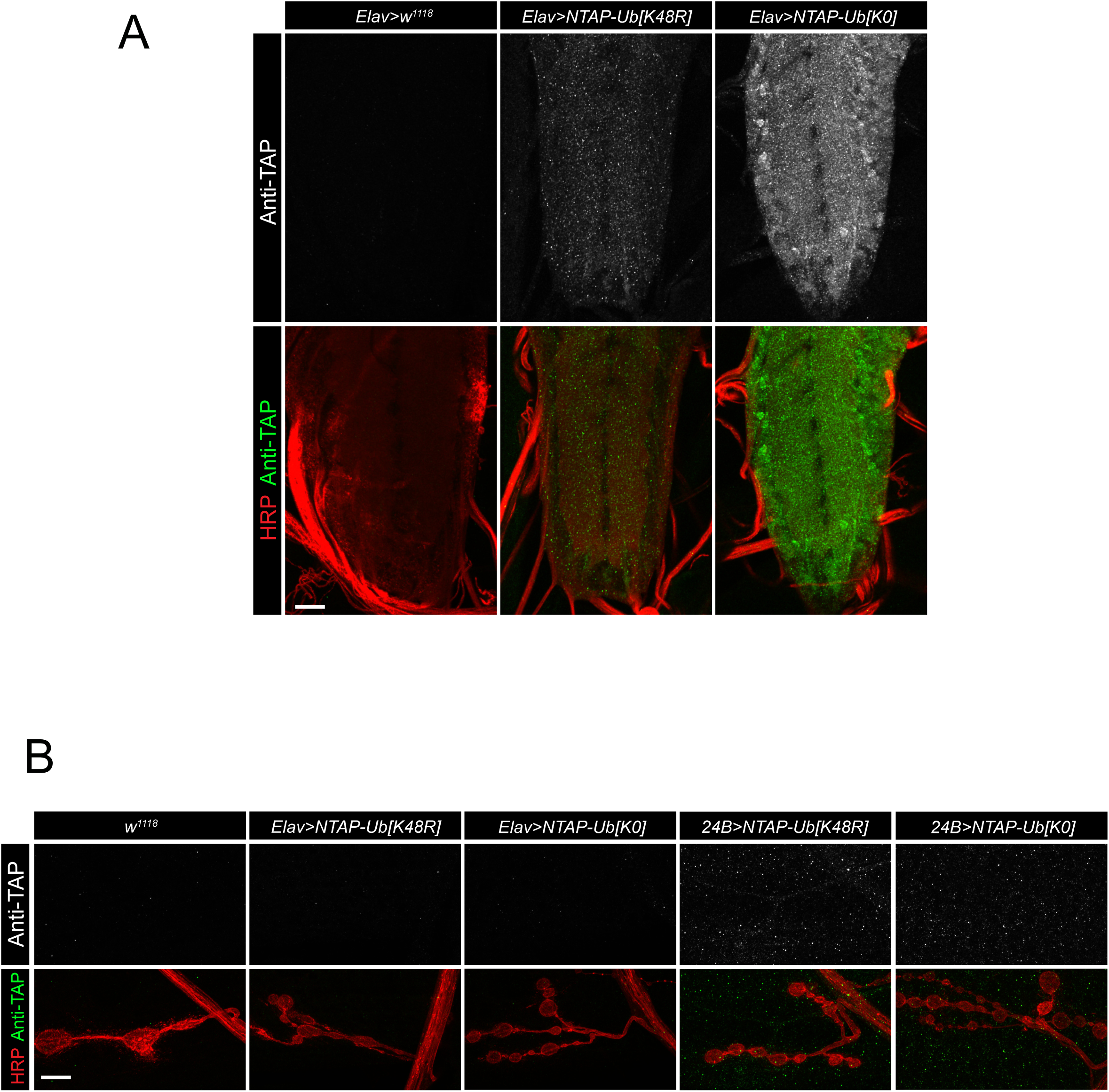
Expression of NTAP-Ub[K48R] and NTAP-Ub[K0] in the larval VNC and NMJs (**A**) Representative confocal images of larval VNCs from control (*Elav>w¹¹¹⁸*) and flies expressing Elav-driven *UAS-NTAP-Ub[K48R]* or *UAS-NTAP-Ub[K0]*. Samples were immunostained with rabbit anti-GFP and goat anti-HRP antibodies. The rabbit anti-GFP antibody was used to detect the TAP tag because it, along with many other rabbit antibodies, cross-reacts with the protein A domain of the TAP tag through its IgG-binding region. All panels show Z-stack projections encompassing the entire VNC. Scale bar: 20 μm. (**B**) Representative confocal images of larval muscle 4 NMJs from control (*w¹¹¹⁸*) and flies expressing *UAS-NTAP-Ub[K48R]* or *UAS-NTAP-Ub[K0]* under the control of Elav-Gal4 or 24B-Gal4 drivers, respectively. Note the absence of NMJ-specific enrichment of TAP-positive staining. Scale bar: 10 μm.

